# Optimization of wheat breeding programs using an evolutionary algorithm achieves enhanced genetic gain through strategic resource allocation

**DOI:** 10.1101/2025.05.01.651521

**Authors:** Azadeh Hassanpour, Antje Rohde, Henner Simianer, Torsten Pook

## Abstract

Plant breeding is a complex process that involves trade-offs among competing breeding objectives and limited resources. Despite the necessity for optimization of breeding program design, the inherent complexity can make optimization challenging. Most often, a predefined set of scenarios is compared or a single parameter of a breeding scheme is assessed in depth. In a previous study, we developed an optimization pipeline, utilizing stochastic simulations and an evolutionary algorithm, suitable for the joint optimization of multiple class and continuous parameters. Here, we assess the applicability of our framework to realistic plant breeding schemes. For this, a wheat line breeding scheme simulated using AlphaSimR and a wheat hybrid breeding scheme simulated using MoBPS were considered. Both schemes were further optimized using our optimization pipeline. When aiming to maximize genetic gain while maintaining a fixed budget, the breeding program design suggested by our optimization pipeline results in 32.6% higher genetic gain compared to a proposed baseline wheat line breeding program. When adjusting the breeding objective to put 20% of the weight on the maintenance of genetic diversity, genetic gain still increased by 4.5% while maintaining 9.1% higher genetic diversity compared to the baseline. Similarly, 4.6%/8.8% higher genetic gains for the male/female part of the hybrid breeding scheme compared to its baseline scenario were obtained. Results highlight the importance of optimizing breeding program design to improve breeding efficiency, with the suggested pipeline offering breeders a powerful framework to refine breeding designs, balance breeding goals, and enhance competitiveness, profitability, and sustainability.

Core Ideas

Evolutionary algorithm can improve breeding design by optimizing many breeding decisions simultaneously

Our framework is compatible with various backend simulators, enabling broad application across platforms

Optimized designs enhance genetic gain and diversity, demonstrating greater overall breeding program efficiency

Genetic gain in wheat line program increased by over 30% compared to a baseline program without additional costs

Plain Language Summary

Modern plant breeding programs must make many complex decisions, such as how many plants to test or where to grow them while staying with tight budgets. These choices are often connected and involve trade-offs between goals like improving genetic gain, maintaining diversity, and reducing costs. In this study, we used an evolutionary algorithm framework to evaluate thousands of breeding program designs and identify the most effective ones. The optimized programs outperformed the respective baseline programs, increasing genetic gain by up to 32% and preserving 9% more genetic diversity - all without additional costs. This flexible framework can support a wide range of breeding decisions and adapt to different program types, providing breeders with a powerful tool to improve outcomes and use resources more efficiently.

## 1 INTRODUCTION

Genetics and breeding technologies are advancing rapidly, driven by innovative techniques and the need for improved crop varieties. These developments enable strategic changes in crop development, which commercial breeders aim to implement efficiently. While the scientific foundation for the quantitative genetics theory and population genetic improvement is established (Falconer & Mackay, 1996), breeding program designs require complex decision-making to address multiple, competing breeding objectives. To meet the needs of a growing global population, expected to reach 9 billion by 2050 (Gerber et al., 2024; Tomlinson, 2013; van Dijk et al., 2021), with significant changes in global temperature (Jaggard et al., 2010), crop production must increase by more than 2% annually to meet future demands (FAO, 2017; Ray et al., 2013), a target that exceeds historical yield growth and underscores the need for highly effective, optimized breeding strategies.

Syngenta reported that developing a soybean cultivar requires making 200 breeding decisions over six years during the variety development process (Byrum et al., 2016). These decisions include important aspects of breeding program design, such as (i) which traits to prioritize (Peng et al., 2014), (ii) how to create new genetic diversity through crossing selected parents (Sebastian Michel et al., 2022), (iii) how to evaluate and select lines across multiple stages and environments (Tolhurst et al., 2019), (iv) and how to advance promising lines for further testing or potential release (Antoine Allier et al., 2020; Asoro et al., 2011; Gorjanc et al., 2017; J. M. Hickey et al., 2014; A. J. Lorenz et al., 2012; Sonja Michel et al., 2017; Windhausen et al., 2012; Zhao et al., 2012). Decisions made at one stage can significantly influence subsequent stages and are often constrained by financial resources (Berry, 2015; Henryon et al., 2014; J. M. Hickey et al., 2017; Simianer et al., 2021). Therefore, breeders must consider not only the technical feasibility of improving certain traits but also the potential market value and long-term impact of these improvements on the germplasm (Brennan & Martin, 2007; Henryon et al., 2014; Simianer et al., 2021).

Over the years, breeders have gained valuable expertise in navigating the complexities of breeding programs, by working on the basis of the breeder’s equation (Falconer & Mackay, 1996; Lush, 1947), by managing breeding schemes effectively within their specific crop and breeding objectives while working with limited resources to balance short-term needs with long-term goals. Certain advancements, such as genomic selection (GS) (Meuwissen et al., 2001), can be used to enable selection in early generations, allowing breeders to make selection decisions without waiting for extensive phenotyping (J.-L. Jannink et al., 2010). Gaynor et al. (Gaynor et al., 2017) proposed an innovative two-part strategy to separate the breeding process into a population improvement component focusing on developing improved germplasm through recurrent GS and a product development component identifying new inbred varieties within conventional breeding designs, which reduced cycle time eightfold compared to a phenotypic selection program, and thereby accelerate genetic gain (Gorjanc et al., 2018). Speed breeding is another technique that can drastically shorten generation times in various crops (Das et al., 2020; Gantovnik et al., 2003; Ghanim et al., 2024; Gudi et al., 2022; L. T. Hickey et al., 2017; Zhang et al., 2017). This method often involves optimizing environmental conditions to accelerate plant growth and development, allowing multiple generations to be produced within a year, and genetic gain can be substantially enhanced beyond two cycles per year (Gaynor et al., 2017; Gorjanc et al., 2018). However, reducing generation intervals increases inbreeding rates and reduces selection accuracy due to omitted phenotyping in multiple generations (Bernardo & Yu, 2007; Gianola & van Kaam, 2008; J.-L. Jannink et al., 2010; Lorenzana & Bernardo, 2009), which can compromise long-term gains.

Given these difficulties and resource constraints inherent in breeding programs (Henryon et al., 2014; Simianer et al., 2021), breeders often exercise caution to depart from long-established breeding practices when considering new approaches (Cobb et al., 2019; Lenaerts et al., 2019; Reynolds et al., 2020). Recently, stochastic simulations have gained popularity as an effective tool to assess these risks, by generating a digital twin of the breeding program on which changes in breeding program design can subsequently be assessed (Faux et al., 2016; Liu et al., 2018; Pook et al., 2020; Sargolzaei & Schenkel, 2009). For the purposes of breeding program optimization, stochastic simulations act as a tool to derive the outcomes of a specific breeding scheme to assess their value towards a predefined breeding target, e.g., the resulting genetic gain. With the use of stochastic simulations, various optimization strategies for breeding programs have been developed to increase genetic gain, many of which focus primarily on the comparison of a set of predefined scenarios or the optimization of individual parameters/components of the breeding program, such as parent selection (Allier et al., 2019; Gorjanc et al., 2018), mating design (Wellmann, 2019; Woolliams et al., 2015), and inbreeding management (Endelman, 2024). Furthermore, stochastic simulations inheritably require high computational power (Pook et al., 2021), which increases exponentially when exploring combinations of parameters (Pook et al., 2025).

Jannink et al. (J.-L. Jannink et al., 2025) and Diot and Iwata (Diot & Iwata, 2022) proposed the use of Bayesian optimization approaches for breeding program optimization. Jannink et al. (J.-L. Jannink et al., 2025) focused on optimizing selection strategies in GS frameworks, presenting improvements in genetic gain by tailoring breeding schemes to specific crop scenarios. Diot and Iwata (Diot & Iwata, 2022) extended this approach and showed the advantage of Bayesian optimization over repeatedly sampling breeding scheme parameters from their prior distributions. Bayesian optimization has shown potential because it can model complex objective functions with fewer evaluations (Wang et al., 2016), making it particularly appealing for constrained optimization tasks. However, the reliance on surrogate models (Frazier, 2018; Shahriari et al., 2016), such as Gaussian processes, limits its scalability and adaptability in high-dimensional spaces or complex breeding schemes. In conclusion, Bayesian optimization is suited for continuous design parameter problems with fewer parameters (Frazier, 2018) while not being suited for class parameters.

In our previous work (Hassanpour et al., 2024), we introduced a novel pipeline based on an evolutionary algorithm (EA) for the optimization of breeding program design with continuous and class design parameters. To reduce the impact of the stochasticity of simulations, kernel regression is used (Hassanpour et al., 2023), and the pipeline is implemented within a parallelized workflow using Snakemake (Mölder et al., 2021). Although the framework is designed to scale well for a high number of parameters, in the context of Hassanpour et al. (Hassanpour et al., 2024) only a simple dairy breeding scheme with three parameters was considered.

Optimizing breeding program design using the EA-based pipeline involves several key steps (Figure 1; Hassanpour et al. (Hassanpour et al., 2024)). First, the breeding problem is stated as an optimization problem, which includes specifying the breeding goals, design parameters, and constraints in the choice of these parameters (Step 0). Next, an initial set of potential parameter settings (corresponding to a specific breeding program design) is generated, providing a starting point for optimization (Step 1). Each parameter setting is then assessed through stochastic simulation, evaluating their performance relative to the objective of the breeding program (e.g., maximizing genetic gain, minimizing economic cost, or preserving genetic diversity) (Step 2). Following this assessment, the most promising parameter settings are selected as "parents" (Step 3), based on which new parameter settings to assess next (“offspring”) are produced (Step 4). The assessment, selection, and generation of new parameter settings are repeated over successive iterations until a stable solution to the objective function is achieved (Step 5). The finally obtained optima are subsequently simulated multiple times to ensure the reliability and consistency of the results (Step 6). For a comprehensive explanation of each step and different techniques for identifying the most promising parameter settings and generating new ones, interested readers can refer to Hassanpour et al. (Hassanpour et al., 2024).

**Figure 1.**
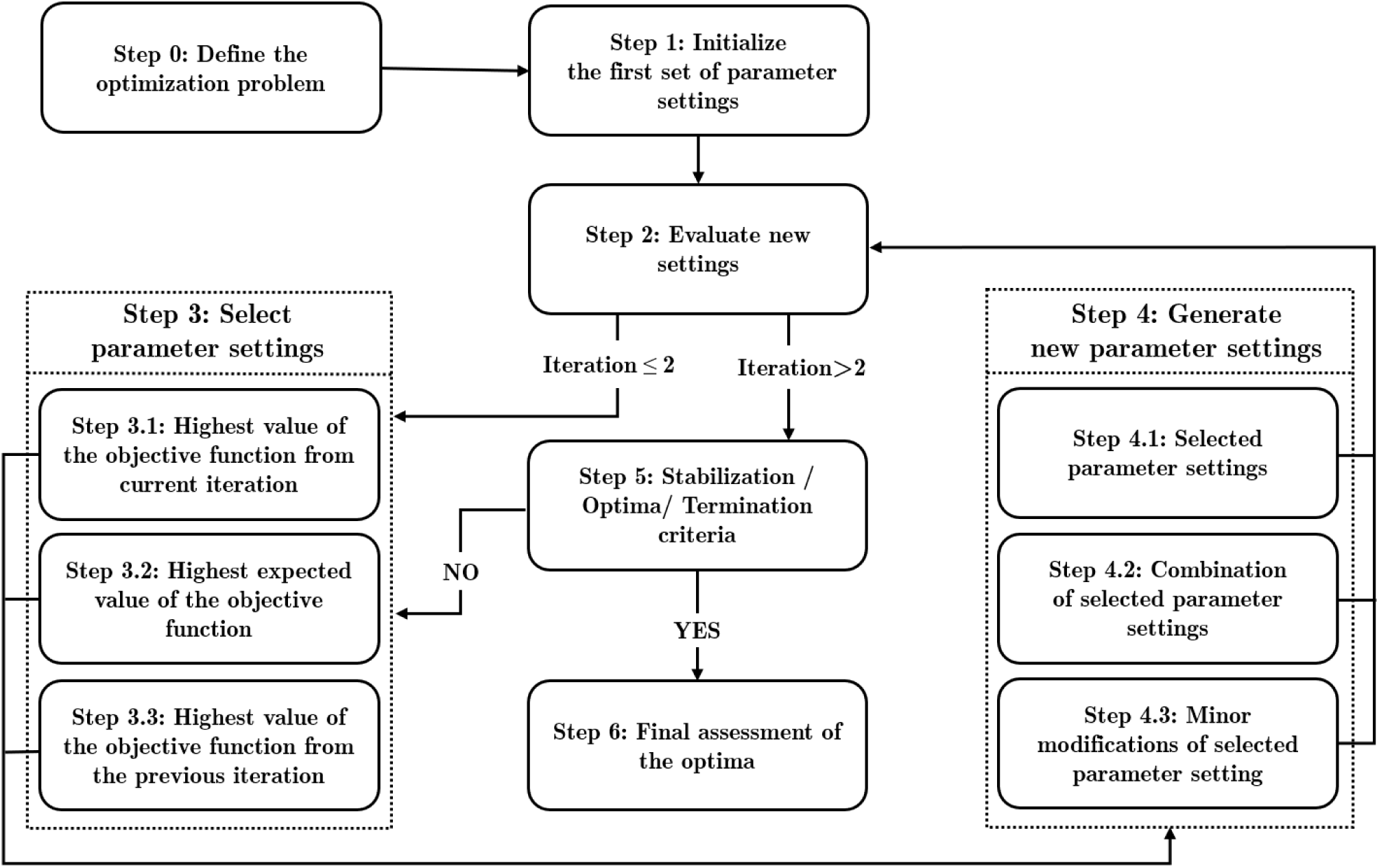
Procedure proposed for optimization via evolutionary algorithm. This figure is taken from Hassanpour et al. (Hassanpour et al., 2024).

Here we show the general applicability of the EA-based optimization framework for breeding program design optimization in plant breeding programs. We apply the EA-based optimization framework to both a wheat line breeding program and a wheat hybrid breeding program, jointly optimizing six and seventeen design parameters, respectively. To further emphasize the framework’s versatility, we employ different backend simulators for the two breeding programs: AlphaSimR (Faux et al., 2016) and MoBPS (Pook et al., 2020). Our optimization focuses on key breeding decisions, including the number of individuals at each stage (cohort size), recycling strategies to select parents for the next breeding cycle, resource allocation for phenotyping and genotyping, and financial investment distribution across program components. Finally, the optimized breeding program design is evaluated and compared against baseline breeding schemes. Subsequently, differences in design are discussed, not only in the context of the given breeding schemes but also providing more tools for a general assessment and optimization of breeding program design through our EA-based optimization framework.

## 2 MATERIALS AND METHODS

In this study, we evaluated two distinct wheat breeding programs using two different software programs to perform stochastic simulations. We selected realistic breeding schemes as baseline scenarios, which serve as effective benchmarks for assessing if and by how much breeding success/gain can be improved by efficient optimization, as conducted through the proposed EA framework.

### 2.1 Optimization of a wheat line breeding program

A wheat line breeding scheme, previously used by Bančič et al. (Bančič et al., 2024) to illustrate the advantages of integrating GS early in the breeding process, is used as a baseline representing current breeding practices. In this study, we consider here the “GS-constrained” scenario as the baseline scenario. A summary of this baseline breeding program, including its sizes and costs, is given in Figure 2 and Table 1. For a detailed description of the simulation methodology, readers are referred to Bančič et al. (Bančič et al., 2024).

**Figure 2.**
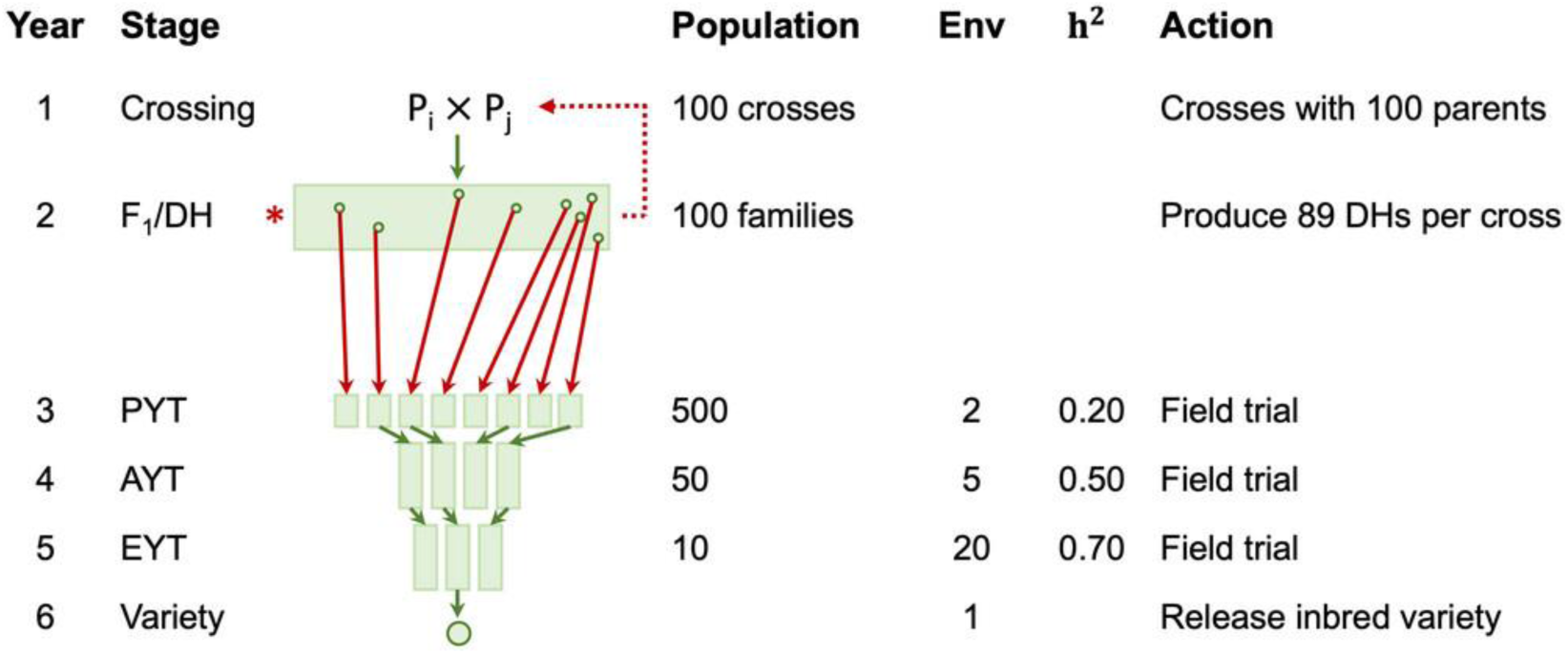
Schematic overview of a wheat line breeding program, highlighting key stages that utilize various cohorts: doubled haploid (DH), preliminary yield trial (PYT), advanced yield trial (AYT), and elite yield trial (EYT). The red dotted line represents the selection of superior parents through recurrent genomic selection (GS), based on their genomic estimated breeding values (GEBV). The figure is taken from Bančič et al. (Bančič et al., 2024) without modification.

**Table 1.** Cost of the baseline wheat line breeding scheme. The table presented here is a simplified version of Table 2 in Bančič et al. (Bančič et al., 2024). For abbreviations, please refer to Figure 2.

| Year | Actions | Cost (\$) | Env | # Units | Cost (\$) |
| --- | --- | --- | --- | --- | --- |
| 1 | Parents | / | / | 50 | / |
| 1 | Cross | 30/cross | / | 100 | 3,000 |
| 2 | Grow F1s | 30/plants | / | 100 | 3,000 |
| 2 | Make DHs | 30/plants | / | 8,900 | 267,000 |
| 2 | Genotype | 15/plants | / | 8,900 | 133,400 |
| 3 | PYT | 20/plot | 5 | 500 | 50,000 |
| 4 | AYT | 50/plot | 15 | 50 | 37,500 |
| 5 | EYT | 50/plot | 20 | 10 | 10,000 |
|  |  |  |  | Total | 504,000 |

In the baseline breeding scheme, each breeding cycle begins with initial crosses in the first year from 50 potential parents. In the second year, the focus is on generating F1/doubled haploid (DH) lines. In the second year, rapid recurrent GS is applied to DH lines, and parents are recycled for the next cycle. From the third to the sixth year, the program advances selected lines to yield trials, involving growing the plants in different environments to evaluate their performance and identify the best-performing lines. At the end of each breeding cycle, the best-performing line will be released as a new variety.

Each cycle will collect the information of all the advanced lines from preliminary yield trial (PYT), advanced yield trial (AYT), and elite yield trial (EYT) to train the GS model with two years of data. In the given baseline breeding scheme, 20% of inbred parents from a total number of 50 parents in each breeding cycle were replaced using newly selected parents, chosen based on their highest genomic estimated breeding values (GEBVs) at the DH stage. The total cost associated with this breeding scheme amounts to 504,000$ in each breeding cycle, of which, in the baseline scenario, 80% is spent in Year 2 for the generation and genotyping of DH lines (Table 1).

For the optimization, we consider here two potential breeding goals, either focusing solely on maximizing genetic gain or a more balanced breeding goal that also aims at maintaining genetic diversity (Bančič et al., 2024). In the following subsections, we describe in more depth the individual steps of applying the EA-based framework for the specific use case.

#### 2.1.1 Step 0: Definition of the optimization problem

The definition of an optimization problem is specific to the application, aiming to identify design parameters that are both impactful but also suitable for change. For the wheat line breeding scheme, we are considering six design parameters:

1. **n_Cross,** the number of crosses among parents to start a breeding cycle
2. **n_DH**, the number of DH lines produced per cross
3. **n_PYT**, the number of entries per preliminary yield trial
4. **n_AYT**, the number of entries per advanced yield trial
5. **n_EYT**, the number of entries per elite yield trial
6. **n_ParentsReplace**, the number of new inbred parents selected each cycle based on GEBVs from the DH stage to replace the oldest inbred parents.

In the following, we are considering the annual budget to be fixed (504,000$) with expenses for each step given in Table 1. A flexible budget could be modelled by integrating a penalty in the objective function (breeding objective) depending on the budget spent. In addition to practical constraints, this results in the following design space:

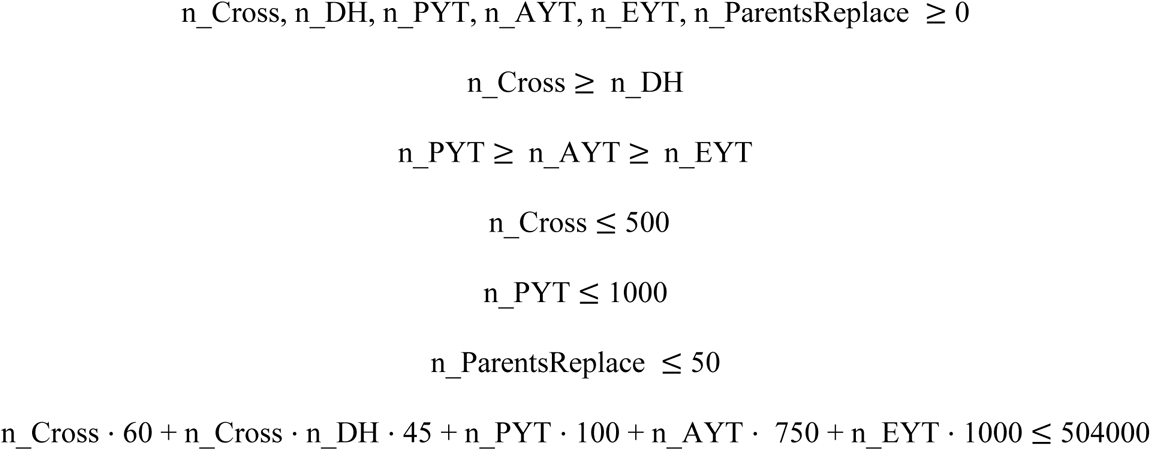

For logistical and practical reasons, the number of crosses (n_Cross) is limited to be at most 500 (e.g., costs associated with making more crosses, managing more field plots, handling additional labels and packets, collecting more data, and conducting more parental checks (Witcombe & Virk, 2001)). At the PYT stage (n_PYT), the maximum field capacity is limited to 1000 lines. The number of parents replaced each year (n_ParentsReplace) cannot exceed the total number of parents (50).

Two distinct breeding objectives were considered for the wheat line breeding program. The first strategy’s objective function aimed to maximize the genetic gain, resulting in a target function *m*:

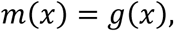

With 𝑔 corresponding to the genetic gain in DH lines after 20 years of breeding, and 𝑥 denotes a potential breeding program design. Note that neither 𝑚 nor 𝑔 can be calculated directly, but simulations will only provide an estimate to derive the expected value (see Step 2). The suggested optimum by the EA will be referred to as "EA gain breeding scheme" throughout the manuscript.

The second breeding objective aims at maintaining a balance between genetic gain and diversity:

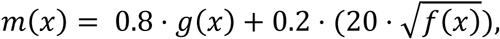

With *f* corresponding to the genetic variance in the breeding values of the DH lines, as used in Bančič et al. (Bančič et al., 2024). As the potential of further genetic improvement is proportional to the genetic standard deviation and not the genetic variance (Falconer & Mackay, 1996), the square root of *f* is used. As *f* and *g* are on different scales, the genetic standard deviation is multiplied by 20 to bring both components on a similar scale. The factor of 20 was chosen based on individual simulations in Bančič et al. (Bančič et al., 2024) showing a standard deviation of approximately 2 units of genetic gain. The objective function is subsequently derived as a composite of the scaled components, in which 80% of the weight is given to genetic gain and 20% to the maintenance of genetic diversity. The suggested optimum by the EA will be referred to as "EA diversity breeding scheme" throughout the manuscript.

#### 2.1.2 Step 1: Initialize the first set of parameter settings

Initial parameter settings (breeding program designs) to consider are generated by random sampling initial values for each design parameter from predefined ranges from a uniform distribution (Table 2). Design parameters may exceed or fall below their initial boundaries during the optimization process if they are not constrained by hard limits from constraints defined in Step 0. After sampling the first set of breeding program designs, all continuous variables (and those that can take a large number of discrete realizations) are scaled to ensure they match the budget, as outlined in our previous work (Hassanpour et al., 2024). In this step, an initial set of 1000 parameter settings is generated.

**Table 2.**
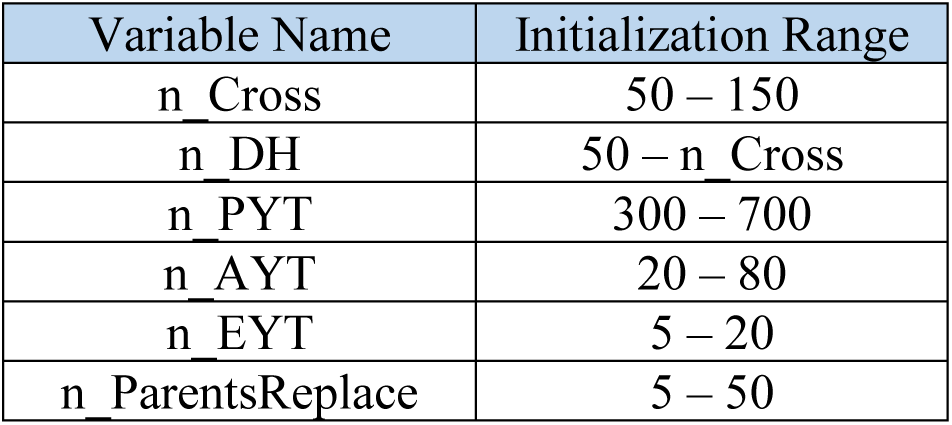
Initial range of design parameters for optimizing the wheat line breeding program. For abbreviations, please refer to Figure 2.

#### 2.1.3 Step 2: Evaluate new settings

To assess the expected outcomes of different breeding program designs, we use stochastic simulation to model each program and evaluate its performance based on the breeding objective. The simulation script was taken from Bančič et al. (Bančič et al., 2024) (available at https://github.com/HighlanderLab/jbancic_alphasimr_plants) and adapted to the EA framework suggested by Hassanpour et al. (Hassanpour et al., 2024) (available at https://github.com/AHassanpour88/Evolutionary_Snakemake/tree/main/script_wheatline).

#### 2.1.4 Step 3: Select parameter settings

We employed the same methodology for selecting optimal parameter settings during the optimization process as outlined in our earlier work in Hassanpour et al. (Hassanpour et al., 2024).

#### 2.1.5 Step 4: Generate new parameter settings

For the generation of new parameter settings, the methodology previously established by Hassanpour et al. (Hassanpour et al., 2024) was adapted to link parameters with each other and allow for ”mutations” in the context of the EA pipeline to affect multiple parameters simultaneously. In this case, this was used as the total number of lines at the end of year two in the breeding scheme results from n_Cross × n_DH instead of a single parameter. For this, ”mutations” are designed not to affect a single design parameter, but act on both n_Cross and n_DH simultaneously to not change the total number of lines generated. Linking parameters here means that if a ”mutation” on one of the parameters occurs, there is a 50% chance that the other parameter is adapted to maintain the same total number of lines generated. To avoid inflation of the total number of ”mutations” in these parameters, ”mutation rates” in these parameters were reduced by 50%. In principle, other types of links between parameters are imaginable, e.g., to increase multiple parameters simultaneously or avoid a high number of ”mutations” overall.

#### 2.1.6 Step 5: Stabilization / Optima/ Termination criteria

As computing times for the wheat line breeding program were low, no early termination criteria were defined. Instead, a fixed number of 150 iterations was performed, followed by visual inspection to determine whether further iterations were necessary.

#### 2.1.7 Step 6: Final assessment of the optima

The suggested optimum after the termination of the iterative optimization is subsequently thoroughly analyzed, and the results are compared with the baseline breeding scheme. For this comparison, genetic gain and diversity at the DH stage per breeding cycle are evaluated based on 100 independent runs for the suggested optimum.

### 2.2 Optimization of a hybrid wheat breeding program

#### 2.2.1 Structure of baseline breeding program

The hybrid wheat breeding program illustrated in Figure 3 is inspired by the two-part strategy proposed by Gaynor et al. (Gaynor et al., 2017), which separates breeding objectives into distinct components. The population improvement component aims to rapidly enhance the population’s average genomic value through recurrent GS, while the product development component aims to identify and develop new hybrid varieties, ensuring that the improved genetic material from the population improvement component is used to create market-ready varieties. In the following, we describe of a typical wheat hybrid breeding program (Figure 3).

**Figure 3.**
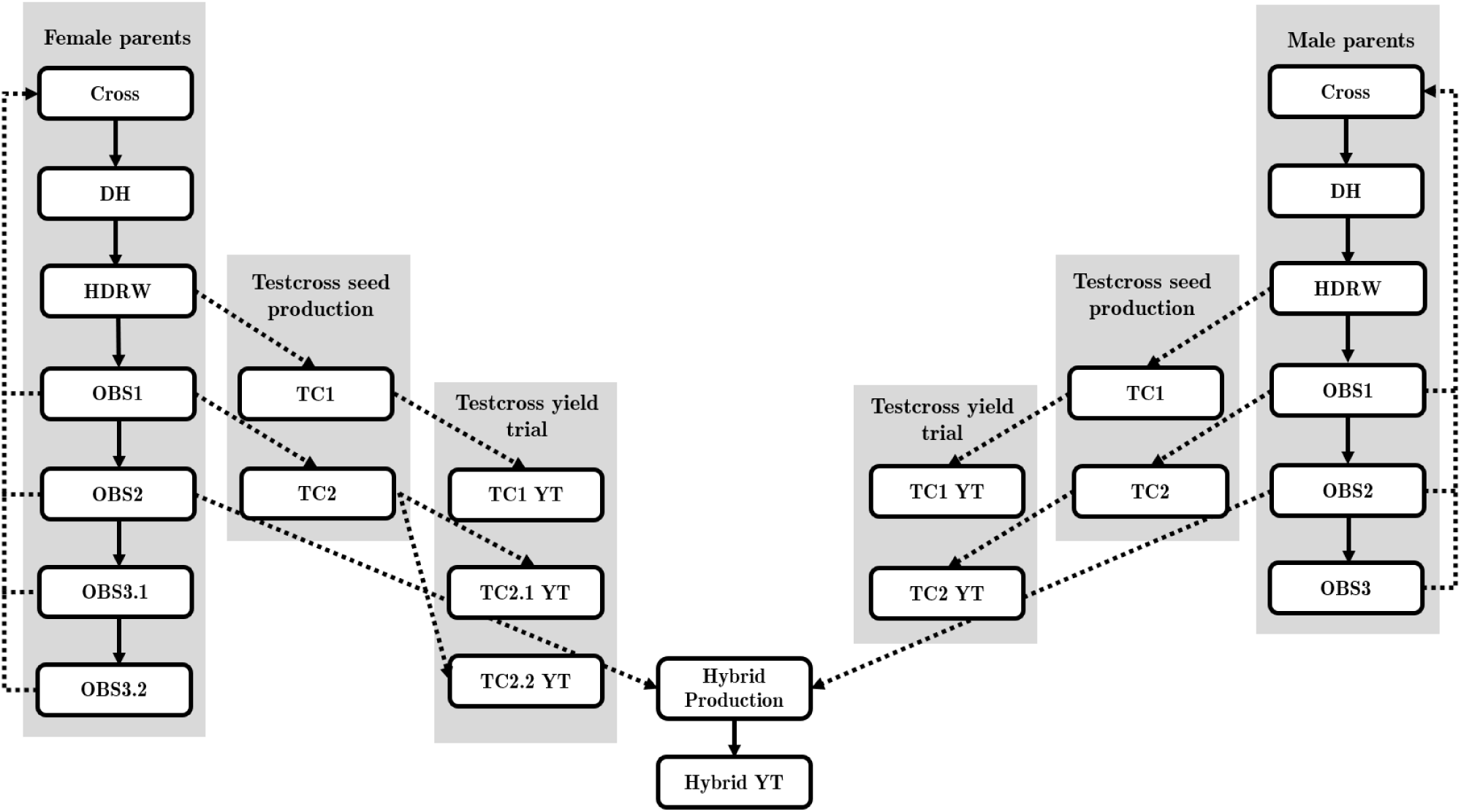
Schematic overview of the hybrid wheat breeding program. The pipeline illustrates the progression through key stages: doubled haploid (DH), headrows (HDRW), observational trials (OBS), testcross seed production (TC), and testcross yield trial (TC YT). An optional repetition of a TC2 trial (TC2.2) is considered to demonstrate its effect.

##### 2.2.1.1 Simulation of the hybrid wheat breeding program

Simulation scripts for this breeding scheme were designed from scratch (https://github.com/AHassanpour88/Evolutionary_Snakemake/tree/main/script_wheathybrid/scripts).

In simulating the hybrid wheat breeding program, we focused on the core genetic aspects and omitted certain practical steps to streamline the model. We disregarded here the incorporation of required elements for pollination control in hybrid production. Further, we did not include seed production stages in the simulation, as these do not directly impact genetic calculations. Similarly, while actual hybrid production was not simulated, its associated costs were incorporated into the overall cost function.

##### 2.2.1.2 eneration of the founder population and traits

The genome was simulated to consist of 21 chromosomes, and assigned a genetic length of 1.43 Morgans, each with 1,000 bi-allelic SNPs. Founder genotypes were generated using the MoBPS function founder.simulation() to build up a basic linkage structure. Three traits were simulated, aiming to mimic grain yield (GY), protein content (PC), and a recessive disease trait (RD). Following Pook et al. (Pook et al., 2025), traits were simulated with separate realizations in male and female parental lines (per se) and hybrids (cross). The correlations between *per se* and cross performances were set to 0.2 for PC and 0.1 for RD. GY was simulated only as a hybrid trait and with a negative correlation to PC (-0.6). The fully considered correlation matrix between traits is given in Table 4.

Furthermore, the phenotyping of individual traits in 10 different environments was considered. Genotype-by-environment interactions (GxE) are simulated by assessing different variants for the different traits depending on the environment. Therefore, for each trait x environment combination, a separate variation of the trait is simulated, resulting in 50 trait combinations when using 5 traits in 10 environments. The required correlation matrix ∑(𝟓𝟎𝐱𝟓𝟎) is generated by combining the correlation matrix between traits∑ (𝟓𝐱𝟓) Kronecker Product:

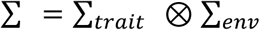

Traits between environments were assumed to be strongly positively correlated, with pairwise correlations between environments sampled to be between 0.6 and 0.8. As pairwise sampling does not necessarily lead to a positive semi-definite matrix, the resulting matrix was subsequently projected in the space of positive semi-definite matrices by setting all negative eigenvalues to 0 (matrix.posdef in MoBPS). As efficient management of GxE is not the primary aim of this study, a relatively simple model was chosen, however, with the availability of real data, more sophisticated simulation approaches like Bančič et al. (Bančič, Gorjanc, & Tolhurst, 2024) exist.

For each subtrait, 300 underlying purely additive and 30 dominant QTLs, with effect sizes drawn from a Gaussian distribution, are simulated with effects shared between traits to obtain the target correlation (see MoBPS Guidelines (Pook et al., 2020); Chapter 15). Traits were assigned weights according to their importance for the desired breeding goal. GY has an index weight of three units per genetic standard deviation (gSD), making it the most important trait, while PC and RD both have an index weight of 1 unit per gSD. QTL effects were scaled to obtain traits with an underlying genetic mean of 100 and a genetic variance of 10 in the first generation.

##### 2.2.1.3 Marker-assisted selection

To mimic the process of marker-assisted selection (MAS), we utilize the underlying true QTL positions, which are known, and phenotypes are regressed against genotypes at a subset of 15% of QTL positions in a linear regression model. To avoid strong selection on individual QTLs, positions used in MAS are resampled for each use of MAS. The choice of 15% is based on empirical sampling to achieve a realistic prediction accuracy in this stage. Trait heritability (h^2^) for each trait and stage is given in Table 2.

##### 2.2.1.4 Outlining the baseline breeding design

At the start (year 1), 100 crosses were created in the female and male parental pool every year (Table 3). Each cross produced 100 F1-derived/DH lines (years 2 and 3).

**Table 3.** Cost and key features of the hybrid wheat baseline breeding scheme with doubled haploid (DH) lines, headrow trials (HDRW), observational trials (OBS), testcross seed production (TC), testcross yield trials (TC YT), replication of testcross yield trials (rep TC2 YT). The considered traits are grain yield (GY), recessive disease (RD), and protein content (PC). The heritability h^2^ values values are not based on formal estimation but are educated guesses derived from practical experience in a real breeding program, considering the fluctuating residual variance observed across various stages of the breeding program. * denotes that 15 hybrids are produced in total, drawing from the identified 30 best female and male DHs.

| Year | Stage | Female population | Male population | Phenotype cost/unit (\$) | Genotype cost/unit (\$) | Environments | Selection GY | $h^2$ GY | Selection RD | $h^2$ RD | Selection PC | $h^2$ PC |
| --- | --- | --- | --- | --- | --- | --- | --- | --- | --- | --- | --- | --- |
| 1 | Cross | 100 | 100 | 30/cross |  |  |  |  |  |  |  |  |
| 2 and 3 | DH | 10000 | 10000 | 30/line |  |  |  |  |  |  |  |  |
| 4 | HDRW | 10000 | 10000 | 10/HDRW | 5 | 1 |  |  | per se | 0,4 |  |  |
| 5 | OBS1 | 500 | 500 | 20/plot | 15 | 2 | pred GCA | 0,5 | per se | 0,4 | pred GCA | 0,5 |
| 5 | TC1 | 500 | 500 | 20/plot |  |  |  |  |  |  |  |  |
| 6 | OBS2 | 300 | 300 | 20/plot |  | 2 |  |  | per se | 0,5 |  |  |
| 6 | TC1 YT | 300 | 300 | 50/plot |  | 2 | GCA | 0,6 | GCA | 0,6 | GCA | 0,6 |
| 6 | TC2 | 300 | 300 | 50/plot |  |  | GCA |  |  |  |  |  |
| 7 | OBS3 | 30 | 30 | 20/plot |  | 2 |  |  | per se | 0,5 | per se | 0,5 |
| 7 | TC2 YT | 30 | 30 | 50/plot |  | 3 | GCA | 0,6 | GCA | 0,6 | GCA | 0,6 |
| 7 | hybrid prod | 15* | 15* | 100/plot |  |  |  |  |  |  |  |  |
| 8 | rep TC2 YT | 20 | - | 50/plot |  | 3 | GCA | 0,6 | GCA | 0,6 | GCA | 0,6 |
| 8 | hybrid YT | 15* | 15* | 50/plot |  | 10 | hybrid | 0,8 | hybrid | 0,8 | hybrid | 0,8 |

**Table 4.** Expanded correlation matrix of traits and the relationships between parental lines (per se) and hybrid (cross) levels for grain yield (GY), protein content (PC), and recessive disease (RD), including their interactions.

|  |  |  |  |  |  |
| --- | --- | --- | --- | --- | --- |
| PC (Per se) | 1 |  |  |  |  |
| RD (Per se) | 0 | 1 |  |  |  |
| GY (Cross) | -0.12 | 0.02 | 1 |  |  |
| PC (Cross) | 0.2 | 0 | - 0.6 | 1 |  |
| RD (Cross) | 0 | 0.1 | 0.2 | 0 | 1 |
|  | PC (Per se) | RD (Per se) | GY (Cross) | PC (Cross) | RD (Cross) |

In year 4, DH lines are planted in headrows, where an initial selection step is performed. As this selection is typically based on visual assessment of plant development and the elimination of undesirable lines based on field observations, it is modeled here as random selection. The remaining lines are subsequently genotyped to apply MAS. Assessments of MAS, along with visual agronomic evaluation, are made in one location. 500 lines are selected after disease evaluation and MAS and enter replicated, multi-location observation trials (OBS1; Table 2).

OBS1 presents the evaluation of DH lines across two environments for the RD trait, while genomic prediction is simultaneously applied to all OBS1 individuals for GY and PC. The training population for GS included all available hybrid data from the past three seasons. The training population is continuously updated in subsequent years by incorporating new yield trial evaluations as they become available. At the same time, the same 500 DH lines/pool are engaged in testcross productions with two testers from the opposite pool to furnish seeds for the subsequent testcross yield trials (TC1, Table 2). The 300 best DH lines are selected based on the *per se* performance of the RD trait and their predicted general combining ability (GCA) for GY and PC traits to advance lines to the first testcross yield trials (TC1 YT, Table 2).

The 300 best DH lines are next tested again in observational trials (OBS2) for RD, and their derived hybrids are tested in testcross hybrid trials for GY, PC, and RD (TC1 YT, Table 2). Concurrently, the same 300 DH lines are engaged in testcross productions with 2 testers from the opposite pool to furnish seeds for the subsequent testcross yield trials (TC2, Table 2). The top 30 parents are selected based on their observed and estimated GCA for all traits within a multi-environment trial (derived from TC1 YT). Additionally, selection considers *per se* performance for RD, derived from OBS2.

The top 30 DH lines are tested further in observational trials (OBS3) for RD and PC, and their derived hybrids are tested in testcross hybrid trials for GY, PC, and RD (TC2 YT, Table 2). Finally, hybrids are produced by making selected combinations with the 30 best females and 30 best males (hybrid prod, Table 2). The top 15 hybrids are selected on GCA for all traits (derived from TC2 YT) and additionally on the *per se* performance for RD and PC (derived from OBS3).

##### 2.2.1.5 Retesting female hybrids

In the female breeding pool, a subset of testcross hybrids undergoes additional evaluation in subsequent testcross yield trials (repTC2 YT, Table 2) to enhance the accuracy of estimated breeding values (EBVs).

##### 2.2.1.6 Recycling of parents

Parents are recycled from multiple stages. 60% of parents are derived from the OBS3 stage (twice evaluated for TC hybrid yield), another 10% come from OBS2 (once evaluated for TC hybrid yield), and a further 10% from OBS1 (only predicted GCA). The remaining 20% of parents consist of new material that, e.g., could come from the characterization of new materials in the pre-breeding program. Newly introduced material is simulated according to the methods suggested by Pook et al. (Pook et al., 2025), by simulated slow repeated recurrent selection in the direction of the expected genetic gain until a similar genetic level to the current breeding population is reached. In this selection process, top-performing candidates with the highest EBVs were allowed to be used approximately five times more often for crosses than the lowest-ranked lines.

##### 2.2.1.7 Filling

In our simulation, we are assuming that the same breeding scheme is executed multiple times, each with a time shift of one year, resulting in all cohorts being generated in the same year, just from different and independently managed breeding schemes that will share information for prediction models and are linked by recycling of lines.

In order to obtain realistic genotypes for all breeding stages, including genetic differences between stages in the base population, an initial filling process similar to Bančič et al. (Bančič et al., 2024) is performed. In the hybrid wheat breeding program, which includes five stages: 1) Cross, 2) DH lines,

1. 3) OBS1, 4) OBS2, and 5) OBS3, the process of filling the pipeline begins by creating and advancing distinct cohorts through each of the stages. Initially, the first cohort is advanced through all five stages, ultimately reaching the final stage (OBS3), making it the oldest cohort. Each subsequent cohort follows, progressing through the stages, with each cohort assigned to a specific stage in the breeding pipeline. The second cohort moves to the fourth stage (OBS2), making it the second oldest, and so on. This continues until all five cohorts are assigned to distinct stages, ensuring that each stage is simultaneously occupied by genetically unique cohorts.

#### 2.2.2 Objectives and parameters for optimization

##### 2.2.2.1 Step 0: Definition of the optimization problem

For the wheat hybrid breeding program, 17 design parameters were identified for subsequent optimization:

1. **rep_TC2_YT f** whether a second testcross yield trial (TC2.2 YT) should be conducted on the female side in year 8 to further select and reduce the number of lines, at the cost of increasing the generation interval
2. **n_Crossf** being the number of crosses on the female side
3. **n_Crossm** being the number of crosses on the male side
4. **n_DHf** being the total number of DH lines produced on the female side
5. **n_DHm** being the total number of DH lines produced on the male side
6. **n_OBS1f** as the number of lines in the observational trial OBS1 produced on the female side
7. **n_OBS1m** as the number of lines in the observational trial OBS1 produced on the male side
8. **n_OBS2f** as the number of lines in the observational trial OBS2 produced on the female side
9. **n_OBS2m** as the number of lines in the observational trial OBS2 produced on the male side
10. **n_OBS3f** as the number of lines in the observational trial OBS3 produced on the female side
11. **n_OBS3m** as the number of lines in the observational trial OBS3 produced on the male side
12. **n_OBS1f_Share** as the proportion of recycling parents selected from the OBS1 female cohort
13. **n_OBS2f_Share** as the proportion of recycling parents selected from the OBS2 female cohort
14. **n_OBS3f_Share** as the proportion of recycling parents selected from the OBS3 female cohort
15. **n_OBS1m_Share** as the proportion of recycling parents selected from the OBS1 male cohort
16. **n_OBS2m_Share** as the proportion of recycling parents selected from the OBS2 male cohort
17. **n_OBS3m_Share** as the proportion of recycling parents selected from the OBS3 male cohort

Hereby, **rep_TC2_YT f** is a binary variable indicating if the yield trial is done (1) or not (0). Cohort sizes are represented as integers, while proportions take numeric values between 0 and 1, making both of these types suitable for treatment as continuous design parameters. Similar to the line breeding program, a fixed annual budget of 2,363,550$ was assumed with expenses linked to each stage given in Table 2. In addition to practical constraints, this results in the following design space:

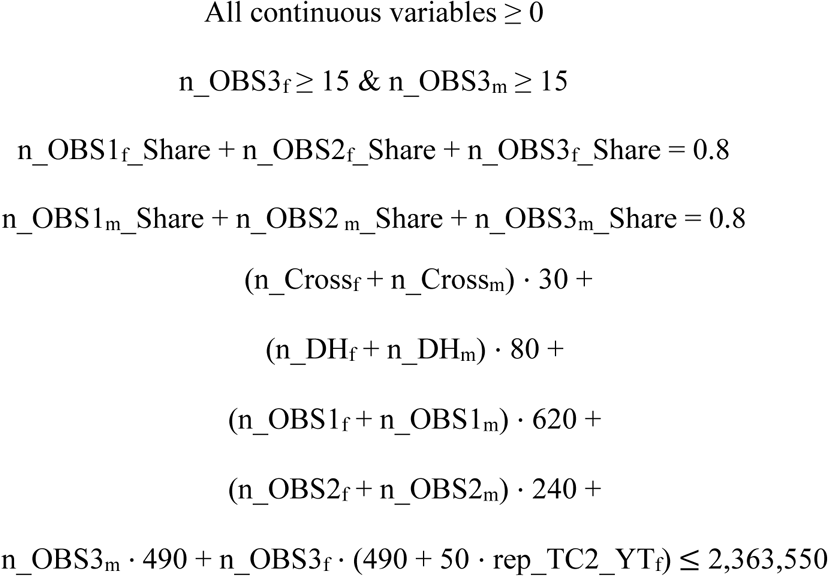

OBS3 is the final step before the product development in the breeding program design begins. To ensure sufficient variation at this stage, the number of n_OBS3f and n_OBS3m is set to a minimum of 15 lines and cannot go below this constraint. The shares of recycled parents selected from the OBS1, OBS2, and OBS3 cohorts were constrained and scaled to sum up to 0.8, ensuring that a constant share of 20% of new genetic material is introduced each year. For this study, we established an objective function to maximize genetic gain achieved over the 20 years of breeding. In many commercial breeding programs, immediate returns are more valuable than future returns. This idea is accounted for by scaling genetic gains by an interest rate (*r*) (Fisher, 1997).

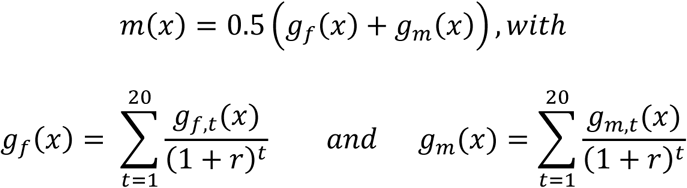

Here 𝑔_𝑓,𝑡_and 𝑔_𝑚,𝑡_represent the expected genetic gain from year 0 to year 𝑡 in the females/males in dependency of the breeding program design parameters 𝑥. In this study, we set *r* = 0.05 (5%) in the objective function, meaning that genetic progress made sooner is weighted more heavily than progress made later.

##### 2.2.2.2 Step 1: Initialize the first set of parameter settings

The initial parameter settings to consider for continuous variables are generated by random sampling for each design parameter from predefined ranges from a uniform distribution (Table 5). rep_TC2_YT f is sampled from a Bernoulli distribution with a probability of 0.5. Similar to the wheat line breeding program, subsequently, scaling on all parameters linked to cost was applied to ensure that all selected breeding program designs matched the target budget. As the overall search space for the hybrid wheat breeding scheme is larger, an initial set of 2000 parameter settings was created to achieve good coverage of the search space.

**Table 5.** Initial range of design parameters for optimizing the hybrid wheat breeding program. For abbreviations, please refer to Table 3.

| Variable Name | Initialization Range |
| --- | --- |
| Binary: Is rep_TC2_YT<sub>f</sub> performed? | 0 (FALSE); 1 (TRUE) |
| n_Cross<sub>f</sub> | 50 – 150 |
| n_Cross<sub>m</sub> | 50 – 150 |
| n_DH<sub>f</sub> | 8000 – 12000 |
| n_DH<sub>m</sub> | 8000 – 12000 |
| n_OBS1<sub>f</sub> | 400 – 600 |
| n_OBS1<sub>m</sub> | 400 – 600 |
| n_OBS2<sub>f</sub> | 200 – 400 |
| n_OBS2<sub>m</sub> | 200 – 400 |
| n_OBS3<sub>f</sub> | 15 – 50 |
| n_OBS3<sub>m</sub> | 15 – 50 |
| n_OBS1<sub>f</sub>_Share | 0 – 0.8 |
| n_OBS2<sub>f</sub>_Share | 0 – 0.8 |
| n_OBS3<sub>f</sub>_Share | 0 – 0.8 |
| n_OBS1<sub>m</sub>_Share | 0 – 0.8 |
| n_OBS2<sub>m</sub>_Share | 0 – 0.8 |
| n_OBS3<sub>m</sub>_Share | 0 – 0.8 |

\

##### 2.2.2.3 Step 2: Evaluate new settings

Expected outcomes of breeding program designs were assessed using the MoBPS stochastic simulator with a simulation script available at:https://github.com/AHassanpour88/Evolutionary_Snakemake/tree/main/script_wheathybrid

As the primary aim of this study was to optimize the population improvement component, product development was simplified in the simulation to exclude final steps of selection that would not be used in recycling steps, as these are assumed to be similar across all scenarios considered.

##### 2.2.2.4 Step 3: Select parameter settings

We employed the same methodology for selecting optimal parameter settings during the optimization process as outlined in our earlier work in Hassanpour et al. (Hassanpour et al., 2024).

##### 2.2.2.5 Step 4: Generate new parameter settings

For the generation of new parameter settings, the methodology previously established in Hassanpour et al. (Hassanpour et al., 2024) was used. Similar to the wheat line breeding scheme, linked parameters were again used. This was done to ensure that the combined shares of recycled parents from the OBS1, OBS2, and OBS3 cohorts always amounted to 80% of the total number of crosses, leading to adaptation of the other two shares in case of a ”mutation”. To prevent an increase in the total number of ”mutations” in linked parameters, the ”mutation rates” for the linked parameters were reduced by 50%.

##### 2.2.2.6 Step 5: Stabilization / Optima/ Termination criteria

Given the high computational cost per simulation and the large number of parameters to optimize, the EA framework for the hybrid wheat breeding scheme was set to run with no specific termination criteria, relying solely on visual assessment.

##### 2.2.2.7 Step 6: Final assessment of the optima

The suggested optimum after the termination of the iterative optimization was subsequently thoroughly analyzed, and its outcomes were compared to those of the baseline breeding scheme based on 100 independent runs for both designs. To evaluate the effectiveness of our framework, the underlying true genomic values for the male and female crosses per breeding cycle were determined. Furthermore, genetic diversity was estimated based on the share of heterozygous markers for the crosses per breeding cycle between scenarios.

### 2.3 Optimization and computing time for simulation

Simulations for the wheat line breeding program, conducted using AlphaSimR simulator Version 1.5.3, required approximately 1 minute and 0.5 GB of peak memory usage per simulation on a single core. Simulations for the hybrid wheat breeding program, using MoBPS Version 1.11.64, required about 40 minutes and 8 GB of peak RAM usage per simulation using 2 cores. All computations were performed on a server cluster with Intel Platinum 9,242 (2X48 core 2.3 GHz) or comparable systems. The EA framework was conducted using the Snakemake workflow management system (version 7.21.0), which distributed individual tasks through a SLURM scheduler.

## 3 RESULTS

### 3.1 Wheat line breeding program

The breeding program designs suggested by the EA pipeline for the wheat line breeding program show substantial differences from the considered baseline (Table 6; Figure S2). For the EA gain breeding scheme, the total number of DHs produced is reduced by 37% to 5,614 DHs. Interestingly, this is primarily achieved by reducing the number of DHs generated per cross (14 instead of 89), while increasing the total number of crosses (401 instead of 100). Subsequently, more resources are allocated to AYT and EYT, increasing their sizes to 230 and 14, respectively (baseline: 50/10). Lastly, in each breeding cycle, 92% of parents were replaced, compared to only 20% in the baseline scenario. Results from 100 simulation replicates demonstrated significant improvements in the optimized breeding schemes compared to the baseline (Figure 6). The EA gain strategy resulted in a 32.6% increase in genetic gain (Figure 6A), with a 73% reduction in genetic variance, compared to a 43% reduction in the baseline (Figure 6B), both relative to year 1.

**Figure 6.**
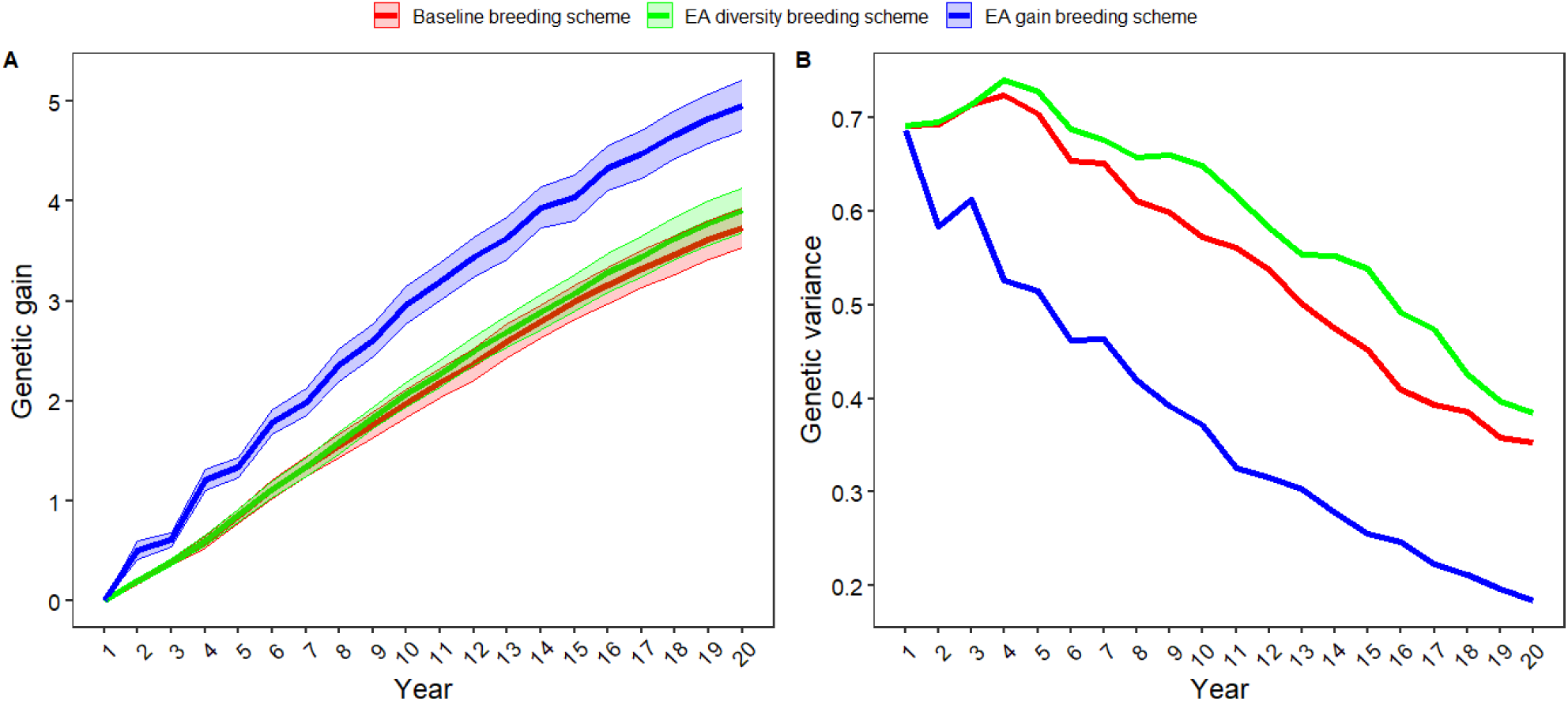
Genetic trends for wheat line breeding program across 100 replicates with **A)** Genetic gain (average increase in breeding values in genetic standard deviations) and **B)** Genetic variance at the doubled haploid (DH) stage. The red line represents the baseline breeding scheme, the blue color represents the EA gain breeding scheme, and the green color represents the EA diversity breeding scheme, all with the same budget.

**Table 6:** Suggested optima for the individual design parameters of the wheat line breeding program design in iteration 150. The baseline values are presented by Bančič et al. (Bančič et al., 2024). For abbreviations, please refer to Figure 2.

| Variable Name | Bancic et al., [63]<br>GS-constrained | Suitable/good parametrization<br>Evolutionary algorithm |  |
| --- | --- | --- | --- |
|  | Baseline breeding<br>scheme | EA gain breeding<br>scheme | EA diversity<br>breeding scheme |
| n_Cross | 100 | 401 | 275 |
| n_DH | 89 | 14 | 32 |
| n_Cross × n_DH | 8900 | 5614 | 8800 |
| n_PYT | 500 | 407 | 677 |
| n_AYT | 50 | 230 | 26 |
| n_EYT | 10 | 14 | 5 |
| n_ParentsReplace | 10 (20%) | 46 (92%) | 10 (20%) |

In contrast, the EA diversity breeding scheme is structurally more similar to the baseline, with a comparable number of DHs produced (8,800). However, the number of DHs per cross is again reduced (32), while the total number of crosses is increased (275). The focus in the yield trial is primarily on PYT, while the number of lines in AYT and EYT is further reduced. Similar to the baseline, only 20% of the parents are replaced in each breeding cycle. Results from the in-depth analysis of the optimum, based on 100 simulations, indicate improvements in both genetic gain (4.5%) (Figure 6A) and remaining genetic diversity (9%) (Figure 6B). Therefore, the breeding scheme is superior in both of these contrasting dimensions, indicating higher efficiency of the breeding program design compared to the baseline.

In terms of overall convergence properties of the evolutionary algorithm, the suggested optima did not improve in their values for the objective function after iteration 50 and 90, respectively, when evaluated across all simulations (Figure S1A). Note that the finally obtained optima for the number of crosses and DHs per cross are both outside of the initially defined search space, which might lead to a longer time till stabilization than when initialized efficiently. The improvement in the objective function was not continuous. In the case of the EA genetic gain scenario, improvements mainly occurred in major jumps in the objective function during two specific iterations (21 and 47). It should be noted that when considering only simulations up to a given iteration (Figure S1A), some variation is observed. This is because the bandwidth used in kernel regression decreases when there are more simulations in the area and simulation outcomes include stochasticity. Even after the overall objective function becomes stable, there is still variation in the proposed optima of individual parameters. However, systematic increases or decreases across several successive iterations are no longer observed. One possible reason is that near the optima, many breeding program designs produce very similar expected outcomes.

### 3.2 Hybrid wheat breeding program

The breeding program design suggested by the EA framework for the hybrid wheat breeding program differs substantially from the baseline (Table 7; Figure S4), with greater emphasis placed on genetic progress achieved earlier in the breeding program. On the female side, the number of crosses is reduced by 76% (24 instead of 100), while the total number of DHs is slightly increased by 11% to 11,052 (baseline: 10,000). The breeding program design places more emphasis on applying higher selection intensity earlier, with the number of female lines in OBS3 reaching the constraint of 15. The generation interval on the female side is slightly increased by recycling lines after OBS1 yield trials (from 60% to 42%) and putting more focus on lines with OBS2 and OBS3 data available (from 10% to 21% and 17%, respectively) compared to the baseline. In contrast, the male side remains close to the baseline, with slightly smaller yield trials suggested across all stages.

**Table 7:** Suggested optima for the individual parameters of the hybrid wheat breeding program design in iteration 75. For abbreviations, please refer to Table 3.

| Variable Name | Baseline breeding scheme | Suitable/good parametrization<br>Evolutionary algorithm |
| --- | --- | --- |
| Binary: Is rep_TC2_YT<sub>f</sub> performed? | TRUE | FALSE |
| n_Cross<sub>f</sub> | 100 | 24 |
| n_Cross<sub>m</sub> | 100 | 87 |
| n_DH<sub>f</sub> | 10,000 | 11,052 |
| n_DH<sub>m</sub> | 10,000 | 9,469 |
| n_OBS1<sub>f</sub> | 500 | 533 |
| n_OBS1<sub>m</sub> | 500 | 448 |
| n_OBS2<sub>f</sub> | 300 | 294 |
| n_OBS2<sub>m</sub> | 300 | 203 |
| n_OBS3<sub>f</sub> | 30 | 15 |
| n_OBS3<sub>m</sub> | 30 | 21 |
| n_OBS1<sub>f</sub>_Share | 0.6 | 0.42 |
| n_OBS2<sub>f</sub>_Share | 0.10 | 0.21 |
| n_OBS3<sub>f</sub>_Share | 0.10 | 0.17 |
| n_OBS1<sub>m</sub>_Share | 0.60 | 0.55 |
| n_OBS2<sub>m</sub>_Share | 0.10 | 0.10 |
| n_OBS3<sub>m</sub>_Share | 0.10 | 0.15 |

Results from 100 simulation replicates indicate substantial improvements in the optimized hybrid breeding scheme (Figure 10). On the female side, the EA scheme results in 8.8% additional genetic gain, and on the male side, the gain is 4.5% higher in year 20 compared to the baseline. For the female side, high additional gain is observed in year 1, due to a reduction in the generation interval (Figure 10A). Subsequent additional genetic gains are similar for the male and female sides and evenly distributed over time (Figure 10B).

**Figure 10.**
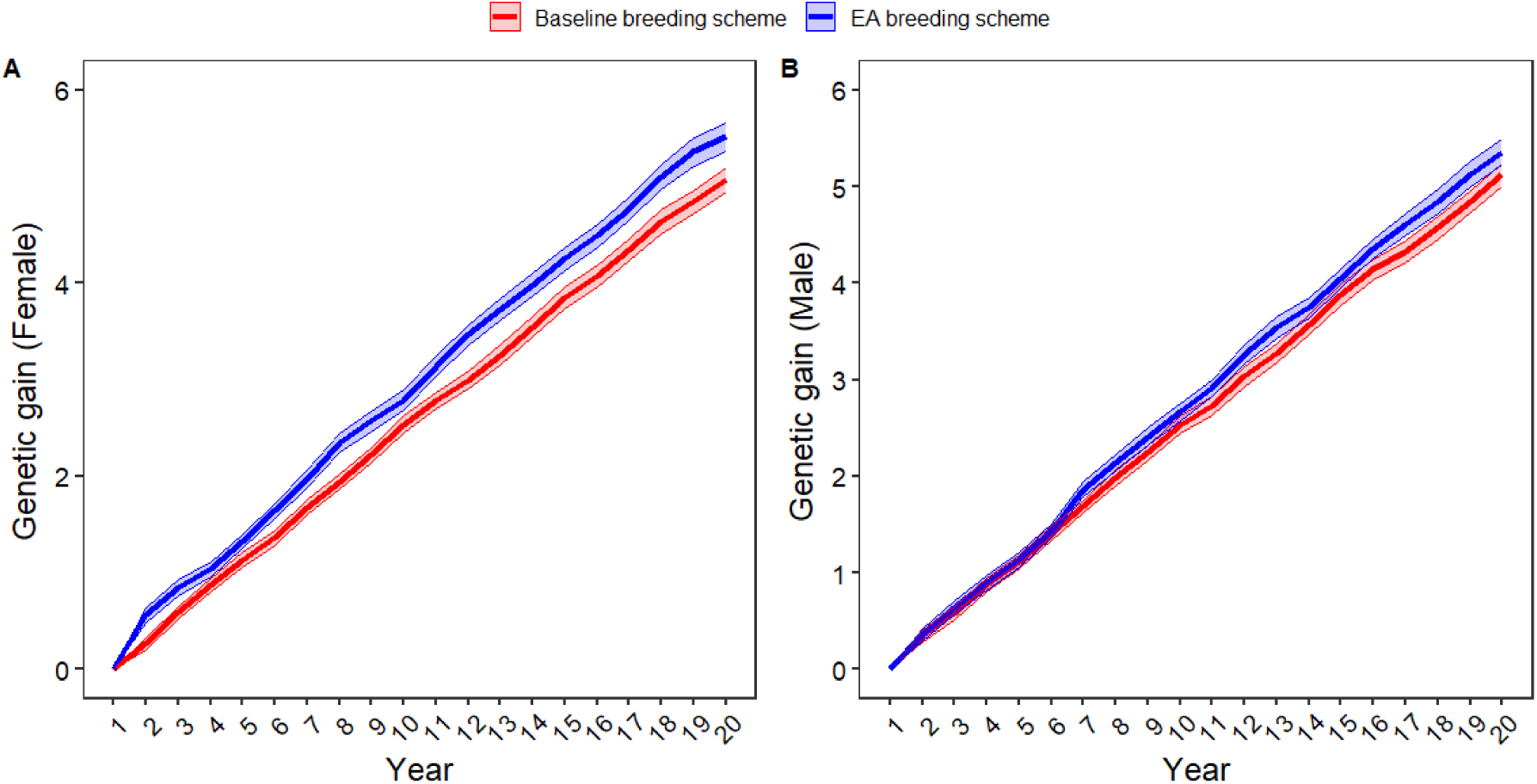
Average increase in breeding values in genetic standard deviations across 100 replicates for **A)** female (n_Crossf) and **B)** male (n_Crossm) breeding population. Blue lines represent the EA breeding scheme using the EA, and red indicates the baseline breeding scheme with the same budget.

Although the baseline showed a slightly higher cumulative gain from OBS2 to OBS3.2 (1.26 vs. 1.22 gSD) on the female side, the small additional gain from OBS3.1 to OBS3.2 in the baseline scheme (0.08 gSD) was offset by more efficient early-stage selection in the EA scheme, with exclusion of the second testcross yield trial (rep_TC2_YTf). A detailed overview of the obtained genetic gains per selection step between scenarios is given in Table 8.

**Table 8.** Comparison of selection intensity (i) and the amount of genetic gain (ΔG) for each step of the breeding programs in both the baseline and EA breeding schemes.

| Step | Baseline<br>Female &<br>Male (i) | EA<br>Female<br>(i) | EA<br>Male<br>(i) | Baseline<br>Female<br>(ΔG) | EA<br>Female<br>(ΔG) | Baseline<br>Male<br>(ΔG) | EA<br>Male<br>(ΔG) |
| --- | --- | --- | --- | --- | --- | --- | --- |
| DH1 → OBS1 | 5% | 4.8% | 7.6% | 0.36 | 0.40 | 0.35 | 0.39 |
| OBS1 → OBS2 | 60% | 55% | 32.7% | 0.32 | 0.35 | 0.31 | 0.45 |
| OBS2→ OBS3.1 | 10% | 5.1% | 6.25% | 1.18 | 1.22 | 1.13 | 1.07 |
| rep_TC2_YT<sub>f</sub><br>OBS3.1 → OBS3.2 | 66% | / | / | 0.08 | / | / | / |
| **Cumulative** | 0.20% | 0.13% | 0.16% | 1.94 | 1.97 | 1.79 | 1.91 |

Over 20 years, both the baseline and EA breeding schemes reduced diversity on both sides (Figure 12). By Year 20, the baseline scheme retained slightly more diversity, with a share of heterozygosity of 29.8%, compared to 28.4% in the EA scheme, compared to 30.5% in year 0. This indicated a much stronger decline in genetic diversity, although overall levels are still high, and gains across the 20 years do not decrease over time. As the EA aimed at optimizing an objective solely focused on increasing genetic gain, no emphasis was put on the maintenance of genetic diversity. However, this could be simply added by modification of the objective function.

**Figure 12.**
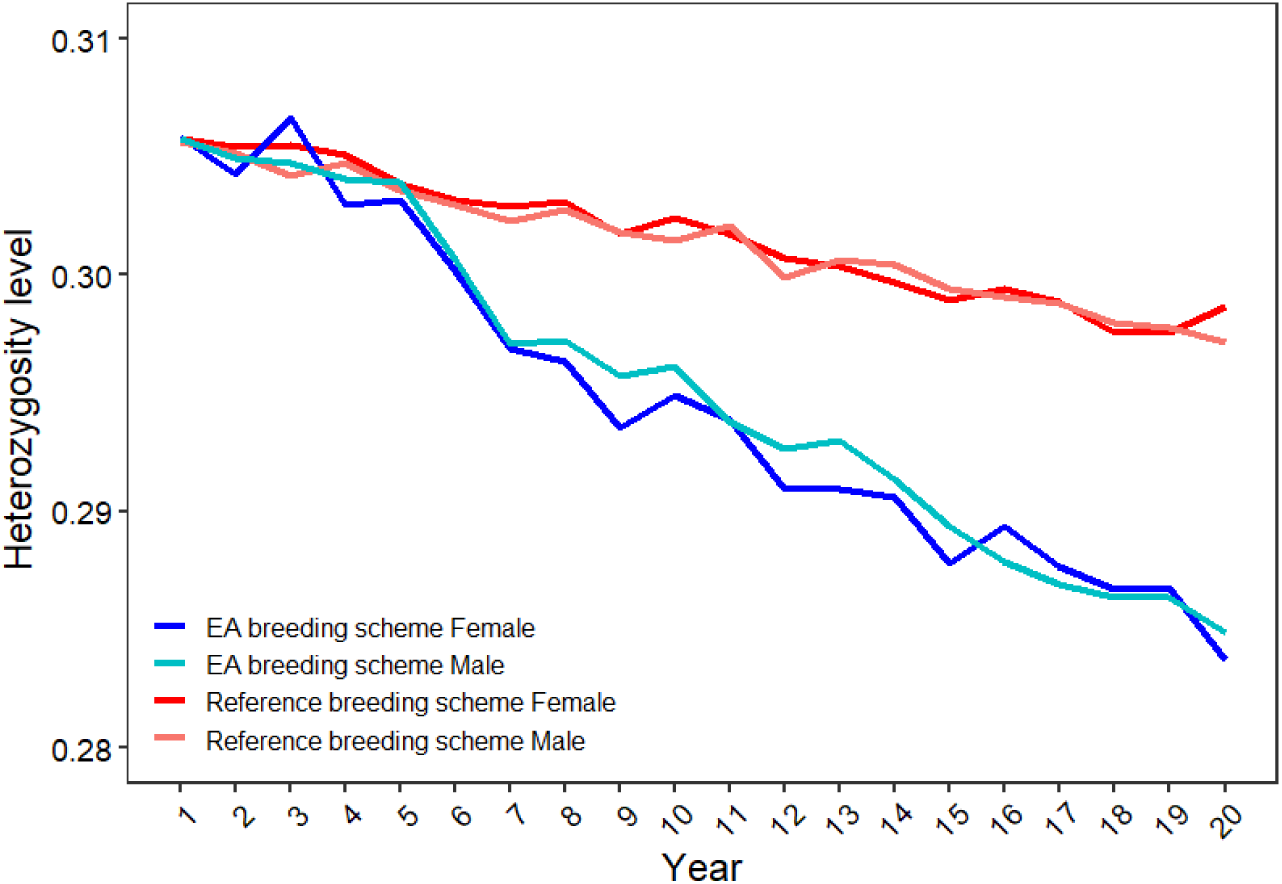
Share of heterozygosity based on the average of 100 replicates for n_Crossf and n_Crossm breeding population. Blue lines represent the EA breeding scheme using the EA, while red indicates the baseline breeding scheme, both of which are implemented with the same budget.

Regarding the overall convergence properties of the EA framework for the hybrid wheat breeding scheme, the proposed optima did not show further improvement in the objective function value after iteration 32, when assessed across all simulations (Figure S3). Some parameters (n_OBS2m_Share (Figure S4O) and n_OBS3m_Share (Figure S4Q) showed considerable change in later generations. However, these changes had only a minor effect on the overall value of the objective function. This suggests that these parameters had limited influence on the final outcomes and could potentially be excluded from the optimization process to obtain faster stabilization. During the optimization process in the hybrid wheat breeding program, the EA quickly identified the exclusion of the second testcross yield trial (rep_TC2_YTf) on the female side as optimal, with the suggested optimum from iteration 14 onwards being unchanged (Figure S4A). The share of selected parameter settings (parents) that include the second testcross quickly drops from 50% in iteration 1 to 10% in iteration 4, and no selection of any of these settings occurs after iteration 24. Note that due to mutations, 1% of the tested parameter settings (population) still include the second testcross in later iterations (Figure S5).

## 4 DISCUSSION

The advancement of highly sophisticated stochastic simulation tools, such as MoBPS (Pook et al., 2020) and AlphaSimR (Faux et al., 2016), has improved the ability to simulate breeding programs that closely mimic real-world scenarios, enabling more effective optimization of breeding strategies. Despite its importance, optimizing breeding program design remains a critical yet often overlooked area of scientific research, as it requires comprehensive knowledge across all components of the breeding pipeline (Kinghorn et al., 2022). As a result, most studies focus on improving individual steps, such as increasing the accuracy of prediction models (Asoro et al., 2011; Zhao et al., 2012; Zhong et al., 2009), enhancing phenotyping and field designs (Araus & Cairns, 2014; Reynolds et al., 2020), or refining selection strategies (Gaynor et al., 2017; Gorjanc et al., 2018). Breeding programs typically operate under strict budgetary and logistical constraints (Aaron J. Lorenz, 2013), making strategic resource allocation essential. Traditional optimization approaches in breeding programs often rely on predefined strategies (Gordillo & Geiger, 2008; Longin et al., 2014) or incremental adjustments (Bančič et al., 2024), which may not fully exploit the potential of optimization methods.

The use of the EA framework (Hassanpour et al., 2024) in this study demonstrates that a systematic optimization approach can improve breeding program design. This improvement was achieved without increasing the budget, through the joint optimization of a high number of independent and dependent design parameters simultaneously, while also integrating the optimization of both class and continuous variables (Eiben & Smit, 2011; Eiben & Smith, 2015; Katoch et al., 2021; Sivanandam & Deepa, 2008). Using the EA framework allows for strategic trade-offs between genetic gain, diversity, and resource allocation (Hassanpour et al., 2024), and the suggested optimized scheme outperformed the baseline in the investigated scenarios. For this study, we used a fixed budget in both breeding programs, but one possible extension is to integrate the budget as an explicit component in the objective function, rather than merely as a constraint to define the search space. This would allow breeders to adjust investments dynamically based on expected returns, allowing them to consider breeding schemes where higher investment would lead to much greater genetic gain (Bančič et al., 2024).

The EA pipeline provides a framework to critically review breeding program design and enhance its efficiency. Exemplary, in the considered baseline wheat line breeding program, approximately 80% of the budget is allocated for genotyping and DH line production. The results from the EA optimizer show high spending in these steps, underscoring their relevance and confirming breeders’ experience and expectations about their importance. Nonetheless, our results also suggest that while a large number of DH lines is necessary to increase the genetic gain by offering more recombination opportunities, the focus should be on selecting a larger number of crosses to avoid discarding potentially favorable ones, while a high number DHs from each cross to chase favorable recombination was not as important (Daetwyler et al., 2015).

Although this conclusion in itself is not novel and has been suggested in previous work (Byrum et al., 2016; Swallow & Wehner, 1989; Wricke & Weber, 1986), the EA framework addresses an important gap by offering a tool that explicitly quantifies trade-offs and supports decision-making. For most critical breeding decisions, valid arguments often exist in multiple directions. This is illustrated by the suggested optimum in the wheat line breeding program, which favored faster replacement of lines to reduce the generation interval, as proposed by Bančič et al. (Bančič et al., 2024). In addition, this study quantified the extent of replacement directly from the objective function. In contrast, the hybrid breeding program prioritized accuracy over generation interval reduction by recycling lines only after more reliable breeding values from multiple environments were available, in line with suggestions from previous works (Bernardo, 2003; Huehn, 2005, 2006). The outcomes of optimization should be interpreted in the given context. When results deviate from expectations, a critical review of the underlying assumptions is essential. For instance, having fewer DH lines per cross for the wheat line breeding program may increase the workload and cost (Witcombe & Virk, 2001), yet this additional labor was not factored into the cost function.

Our study further emphasizes the importance of breeding program design by showing that optimization in the wheat line breeding program increased genetic gains by 30%. Thereby, providing higher additional genetic gains than the integration of GS early in the breeding process (20%), as estimated by Bančič et al. (Bančič et al., 2024) for the same breeding scheme. This highlights that effective design is crucial.

The breeding scheme used for the hybrid wheat program included multiple traits, making the optimization more realistic and better aligned with practical breeding objectives. Even for this design, the results of the EA suggest an improvement of more than 5%. This is particularly appealing when considering that additional gains come without any additional financial investment, providing breeders with a tool to maximize their profitability.

The EA framework offers a flexible optimization tool for addressing general questions in breeding program design, such as optimizing cohort sizes, deciding whether specific yield trials are necessary, or adjusting recycling intervals. Similarly, more focused research questions targeting individual components of a breeding scheme can also be explored with the same approach, but in greater detail. Pook et al. (Pook et al., 2024) used the EA framework to improve selection by comparing and combining various strategies to better account for genetic diversity and long-term genetic gain. In the context of plant breeding, we could particularly envision the use for effective management of GxE by examining how to best allocate resources, such as years, seasons, locations, and replicates, or limited field plots, considering the cost-effectiveness of adding extra replicates compared to additional locations or years (Hanson & Brim, 1963; Sprague & Federer, 1951; Swallow & Wehner, 1989; Wricke & Weber, 1986; Zhou et al., 2011).

For the hybrid wheat breeding program, running 24,600 simulations over 75 iterations, with each simulation taking 40 minutes, results in a computational cost of approximately €650 (based on a cloud computing rate of €0.012 per CPU core hour and €0.012 per 6 GB of memory per hour https://hpc.ut.ee/pricing/calculate-costs), while all other steps of the EA had negligible computational demand. Compared to the total budget typically spent in a real breeding program, this cost is minimal and highlights the advantage of using EA pipeline for efficient and affordable optimization. It also underscores the importance of having efficient simulation software, as well as the ability of the EA to reduce the total number of simulations needed to reach a good solution.

Based on our analysis of the algorithm’s performance across different problems the overall design of the EA pipeline has proven effective across various optimization problems in breeding program design, as demonstrated in this study for the wheat line breeding program, the hybrid wheat breeding program, and in our previous study (Hassanpour et al., 2024).In this study, we extended the previously suggested EA framework (Hassanpour et al., 2024) by introducing linked parameters in the generation of new parameter settings to test and thereby obtain better-suited new candidates and reduce the overall number of simulations needed. Another critical factor for improving EA efficiency is the careful definition of appropriate parameter bounds during initialization (David E. Goldberg, 1989; Goldberg, 1989). A narrow search space will most likely lead to faster convergence, while broader bounds increase the likelihood of identifying globally optimal or innovative breeding program designs. That said, the potential innovativeness of a suggested optimum is ultimately constrained by the design of the simulation script itself, which is user-defined and non-generative in nature.

### 4.1 Conclusion

This study demonstrates the successful application of a combination of stochastic simulation and an evolutionary algorithm for optimization as a powerful tool to support breeders in designing more effective breeding programs. The framework is highly versatile, capable of addressing complex breeding scenarios with numerous design parameters, including both continuous and class variables, and has been successfully applied across line and hybrid plant breeding programs using various stochastic simulators. Importantly, the EA framework is not intended to replace breeder expertise but serves as a valuable decision-support tool that identifies opportunities for optimization within a breeding program. By systematically analyzing a wide range of variables and testing different design scenarios, it uncovers potential improvements that might be overlooked through conventional methods. This enables breeders to make more informed decisions, refining strategies to better align with their program objectives. By refining existing strategies through efficient resource reallocation, breeding programs can better navigate inherent trade-offs, such as investing in phenotyping versus genotyping or choosing between evaluating more candidate lines or focusing on fewer in greater detail.

## Data availability

The presented evolutionary framework is patent pending under application numbers EP24164947.4 and EP24188636.5. Patent applicants are BASF Agricultural Solutions Seed US LLC and Georg- August-Universität Göttingen. Inventors are Torsten Pook, Azadeh Hassanpour, Johannes Geibel, and Antje Rohde. Academic, noncommercial use is possible under a public license, with all examples supported with scripts on the GitHub repository details given at: https://github.com/AHassanpour88/Evolutionary_Snakemake/blob/main/License.md.

## Acknowledgments

The authors acknowledge the computational support from the Scientific Compute Cluster at GWDG, the joint data center of the Max Planck Society for the Advancement of Science (MPG), and the University of Goettingen. We acknowledge support by the Open Access Publication Funds of the Goettingen University. We thank the BASF Agricultural Solutions Biometrics team in Gent for valuable discussions and insights that contributed to this study.

## ABBREVIATIONS

AYT: advanced yield trial
DH: doubled haploid;
EA: evolutionary algorithm
EBVs: estimated breeding values
EYT: elite yield trial;
GCA: general combining ability;
GEBVs: genomic estimated breeding values
GS: genomic selection;
GxE: genotype-by-environment interactions
GY: grain yield;
HDRW: headrows;
MAS: marker-assisted selection
OBS: observational trials;
PC: protein content;
PYT: preliminary yield trial
RD: recessive disease trait
TC: testcross seed production
TC YT: testcross yield trial.

