## Supplementary Material for "Optimization of wheat breeding programs using an evolutionary algorithm achieves enhanced genetic gain through strategic resource allocation"

### Supplementary files

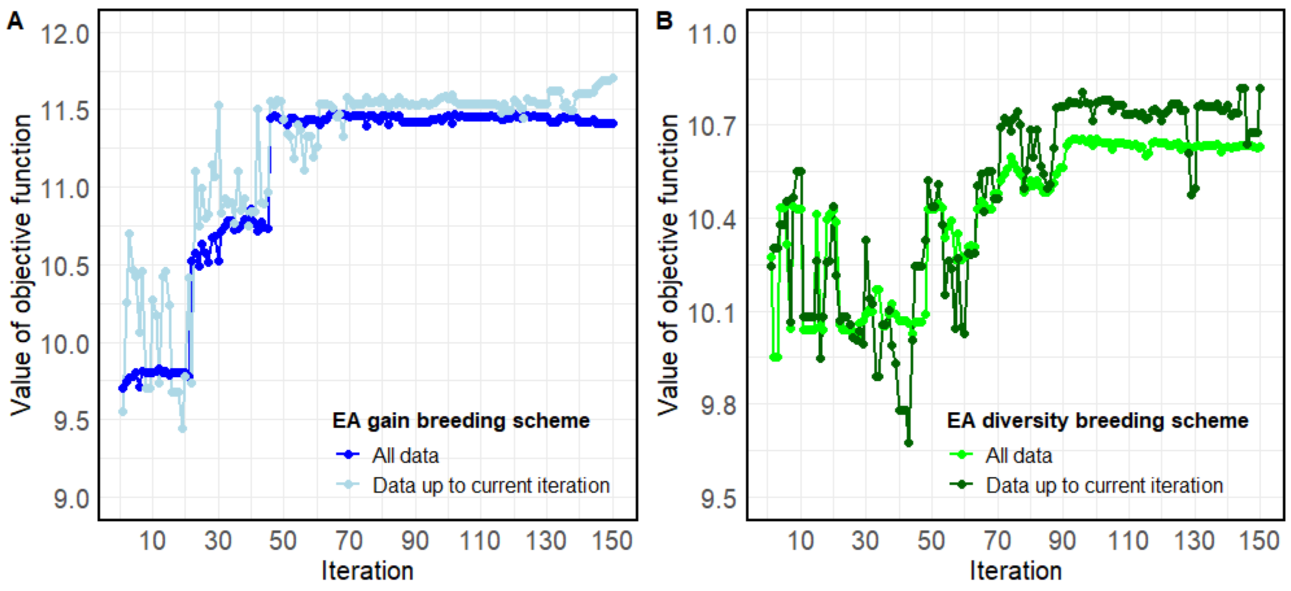


Figure S1. Performance of the suggested optima for the wheat line breeding program after each iteration was assessed using kernel regression for **A)** the EA gain breeding scheme and **B)** the EA diversity breeding scheme.

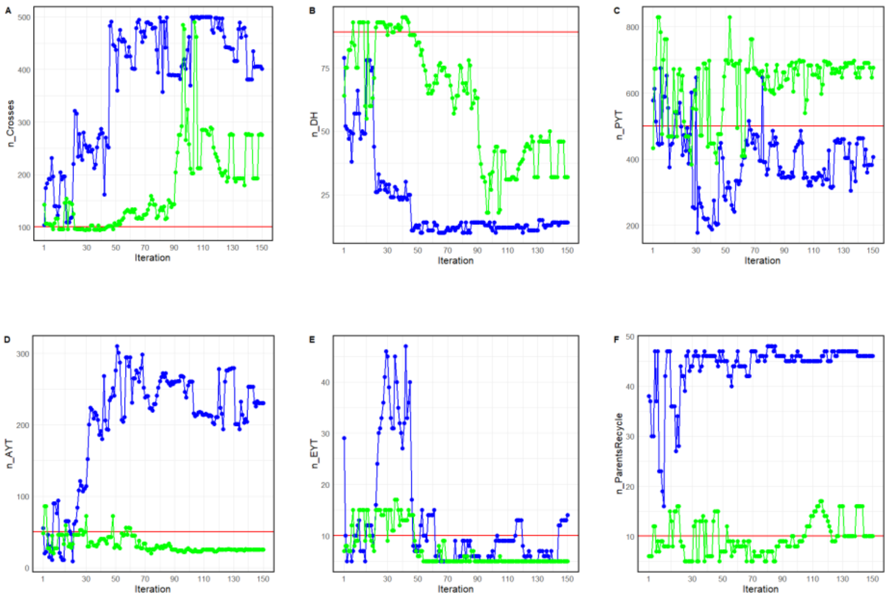


Figure S2. Suggested optima for the individual parameters of the wheat line breeding program design for number of **A)** crosses, **B)** DH lines produced per cross, **C)** entries per PYT, **D)** entries per AYT, **E)** entries per EYT, and **F)** new inbred parents to replace the oldest inbred parents. The red horizontal line represents the reference scenario presented by Bančič et al. [61]. The blue lines indicate the EA gain breeding scheme, and the green lines represent the EA diversity breeding.

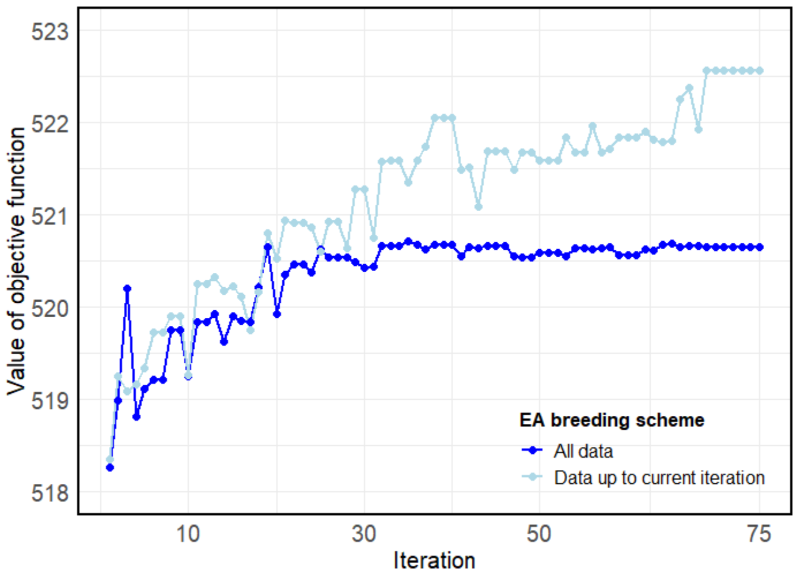


Figure S3. Performance of the suggested optima for the hybrid wheat breeding program after each iteration was assessed using kernel regression.

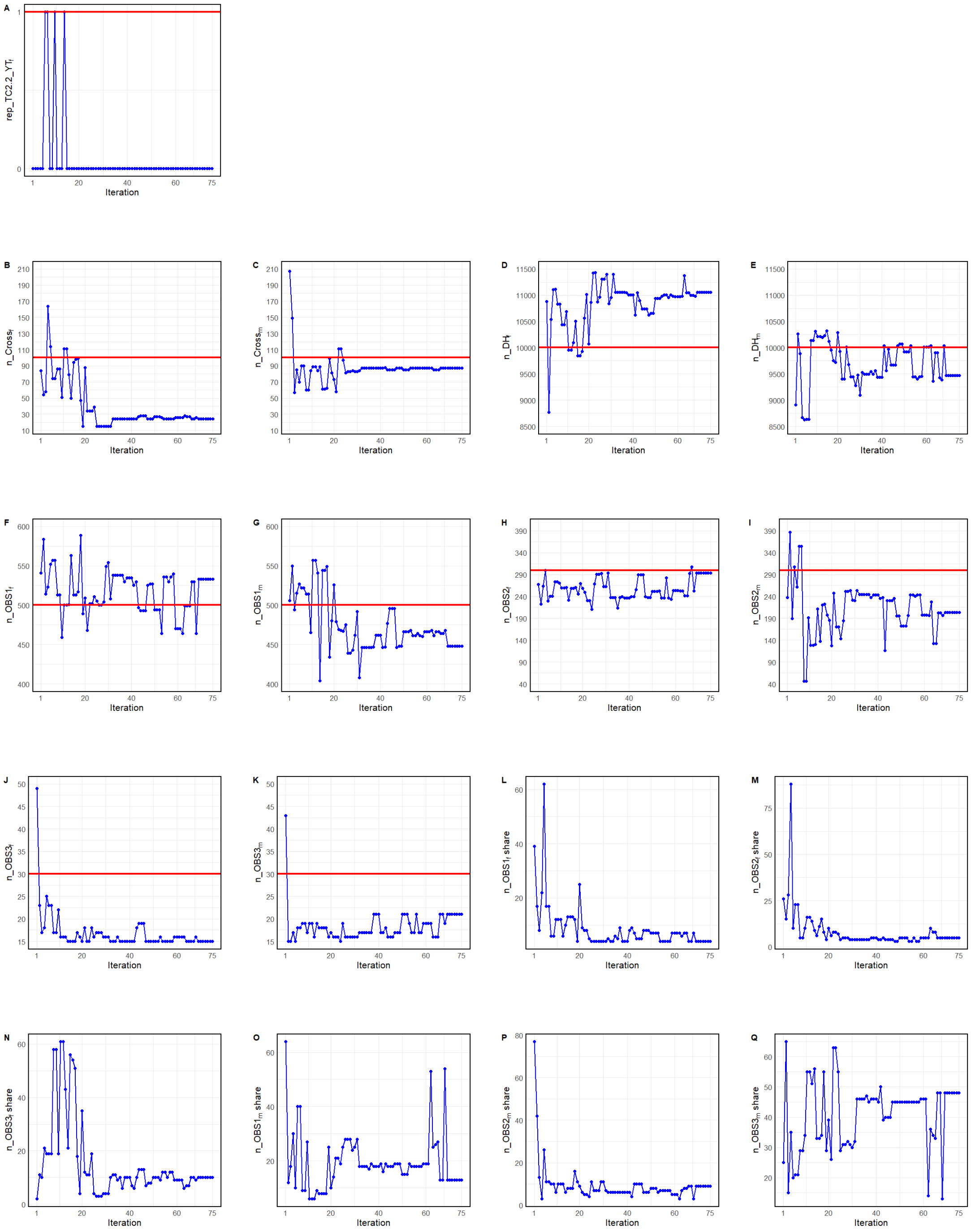


Figure S4. Suggested optima for the individual parameters of the hybrid wheat breeding program design with **A)** rep_TC2_YT $_f$, **B)** n_Cross $_f$, **C)** n_Cross $_m$, **D)** n_DH $_f$, **E)** n_DH $_m$, **F)** n_OBS1$_f$, **G)** n_OBS1$_m$, **H)** n_OBS2$_f$, **I)** n_OBS2$_m$, **J)** n_OBS3$_f$, **K)** n_OBS3$_m$, **L)** n_OBS1$_f$_Share, **M)** n_OBS2$_f$_Share, **N)** n_OBS3$_f$_Share, **O)** n_OBS2$_m$_Share, **P)** n_OBS2$_m$_Share, **Q)** n_OBS3$_m$_Share. The red line represents the baseline breeding scheme, while the blue lines indicate the EA breeding scheme using the evolutionary algorithm.

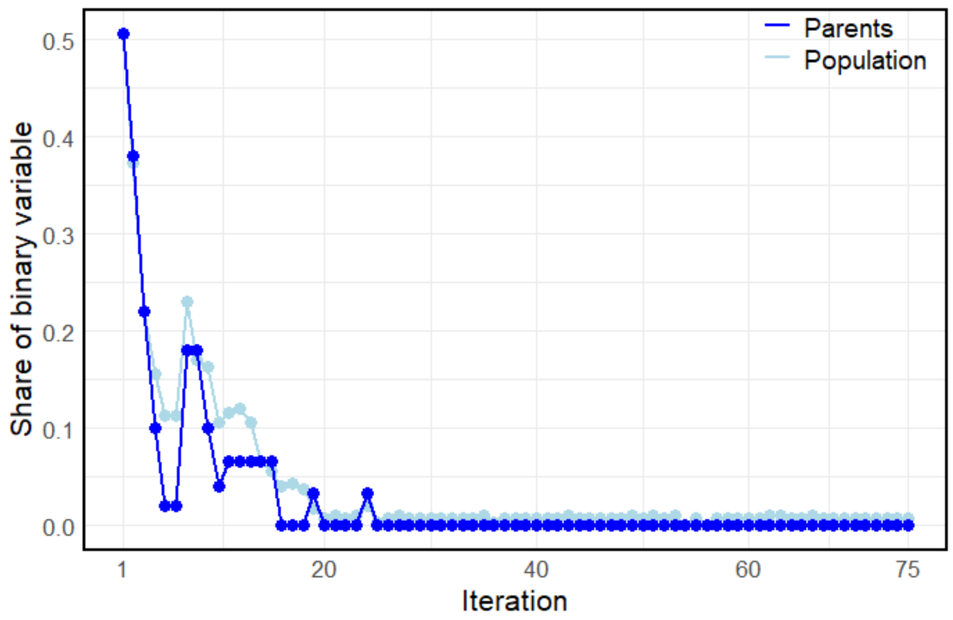


Figure S5. Share of the binary variable to perform replication of the testcross yield trial on the female side rep_TC2_YT$_f$ in year 8 in each iteration for the optimized hybrid wheat breeding program, with blue being the share of parents and lightblue being the share of the population.
